# Organised streaming of cerebrospinal fluid in the human third ventricle

**DOI:** 10.64898/2026.11.03.763888

**Authors:** Pengcheng Zhao, Luke McTavish, Christoph Bruecker

## Abstract

Quasi-periodic pressure pulses associated with cardiac and respiratory activity generate reciprocating cerebrospinal fluid (CSF) motion in the human ventricles. Regular waves dominated by the cardiac cycle are overlaid with cycles of low-frequency respiratory waves, the latter to become stronger in forced breathing or even dominated by large successive flushes in NREM sleep. The present study investigates how such waves contribute to residual streaming within the human ventricles. Transient CFD simulations were performed in the third ventricle at varying parameters of frequency, stroke volume and wave-form asymmetry. Increasing the aqueduct peak Reynolds-number, either by small stroke volume and high frequency or large stroke volume at low frequency, transfers the flow from a reversible flow pattern to the formation of clear preferential pathways, alternating between the in- and outflow cycle. The resulting permanent streaming flux across the aqueduct relative to the mean oscillatory flow-rate, i.e. the dimensionless streaming ratio SR follows a rational saturation relation, *SR ~ Re*_Aq,peak_*/*(*C* + *Re*_Aq,peak_), demonstrating that increasing the peak aqueduct Reynolds number yields diminishing returns and bounds the relative contribution of residual inter-ventricular streaming. These findings demonstrate differences in hydrodynamic organisation in the third ventricle and highlight the importance of this organisation for intraventricular fluid exchange.

## Introduction

Cerebrospinal fluid (CSF) provides mechanical protection for the brain and participates in fluid and solute exchange [1, 2]. Within the ventricles, the net flow associated with CSF production and drainage is accompanied by reciprocating fluid motion driven by cardiac pressure pulsation and respiration [3, 4]. It was observed that this part of reciprocating fluid flow (or bi-phasic flow) is typically about an order of magnitude greater than the net volume displaced in each cycle [5]. The stroke volume, the tidal fluid bulk moved caudal and rostral through the cerebral aqueduct of Sylvius (AoS) in each cycle, has been reported for healthy adults in the normal range from 1.9 *µ*L up to 33 *µ*L, with the average sitting around 17 *µ*L as established by Abbey et al. [6]. Cardiac-gated MRI measurements characterise the waveform of CSF motion as skewed harmonic profile at a dominant frequency of about 1Hz [5], [7]. Modern phase-contrast MRI can provide a fuller picture of the waveform, indicating that breathing is contributing to a larger percentage of the reciprocating flow. Under deep or sleep-augmented respiratory drive, the dominant oscillatory cycle shifts to the respiratory frequency (*~* 0.25 Hz, *T ≈* 4 s), amplifying stroke volumes. Dreha-Kulaczewski et al., (2017) reported in their results (Table 1) from real-time PC-MRI measurements that the aqueduct stroke volume surges up to 150–500 *µ*L (0.15–0.50 mL)—which is 3 to 10 times larger than the 30–50 *µ*L resting cardiac stroke volume (comparing healthy forced breathing vs. quiet breathing). The respiratory macro-wave exhibits a pronounced temporal and amplitude asymmetry, pairing a rapid, steep cranio-caudal surge during active inspiration with a prolonged, blunted caudo-cranial return during passive expiration [8]. Within this respiration-dominated regime, the 0.25 Hz waveform governs bulk advective transport, whereas the continuous 1 Hz cardiac pulsation is reduced to a secondary, low-amplitude ripple [9, 10]. This is especially reported for the situation of forced breathing [4, 11]. Details of the CSF flow motions in sleep still remain obscured due to practical limitations of keeping the cohort asleep in the scanner. Recent studies utilizing ultrafast MR Encephalography (MREG) combined with optical flow analysis [12] have tracked how the 3D velocity dynamics of CSF change the moment a human transitions from wakefulness to Non-REM sleep. They confirmed that sleep induces a massive dynamic shift in the flow speed. The velocity of respiratory pulsations increases by more than 20% compared to the awake state.

**Table 1.**
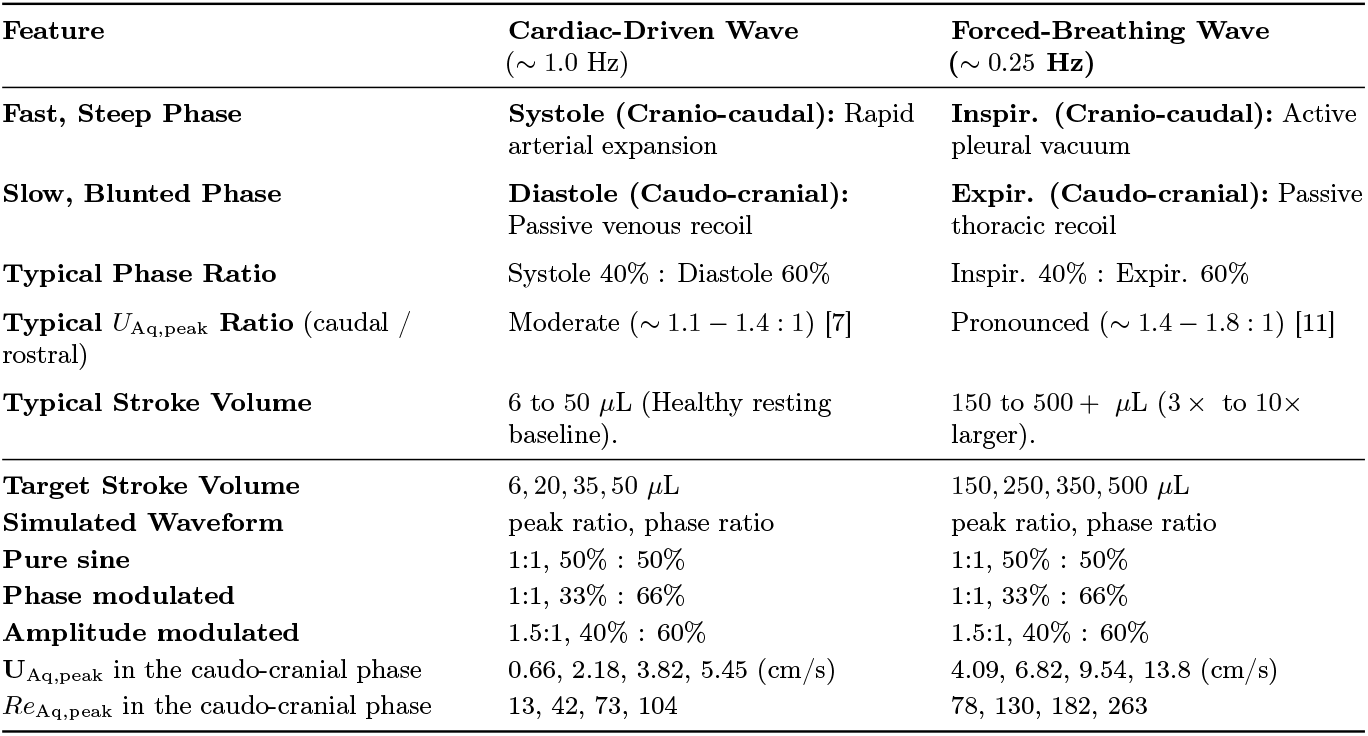
Simulated kinematic and physiological features for cardiac-driven and forced-respiratory driven CSF flow in the cerebral aqueduct.

| **Feature** | **Cardiac-Driven Wave**<br>(∼ 1.0 Hz) | **Forced-Breathing Wave**<br>**(∼ 0.25 Hz)** |
|---|---|---|
| **Fast, Steep Phase** | **Systole (Cranio-caudal):** Rapid arterial expansion | **Inspir. (Cranio-caudal):** Active pleural vacuum |
| **Slow, Blunted Phase** | **Diastole (Caudo-cranial):** Passive venous recoil | **Expir. (Caudo-cranial):** Passive thoracic recoil |
| **Typical Phase Ratio** | Systole 40% : Diastole 60% | Inspir. 40% : Expir. 60% |
| **Typical** *U*<sub>Aq,peak</sub> **Ratio** (caudal / rostral) | Moderate (∼ 1.1 − 1.4 : 1) [7] | Pronounced (∼ 1.4 − 1.8 : 1) [11] |
| **Typical Stroke Volume** | 6 to 50 µL (Healthy resting baseline). | 150 to 500 + µL (3 × to 10× larger). |
| **Target Stroke Volume** | 6, 20, 35, 50 µL | 150, 250, 350, 500 µL |
| **Simulated Waveform** | peak ratio, phase ratio | peak ratio, phase ratio |
| **Pure sine** | 1:1, 50% : 50% | 1:1, 50% : 50% |
| **Phase modulated** | 1:1, 33% : 66% | 1:1, 33% : 66% |
| **Amplitude modulated** | 1.5:1, 40% : 60% | 1.5:1, 40% : 60% |
| U<sub>Aq,peak</sub> in the caudo-cranial phase | 0.66, 2.18, 3.82, 5.45 (cm/s) | 4.09, 6.82, 9.54, 13.8 (cm/s) |
| *Re*<sub>Aq,peak</sub> in the caudo-cranial phase | 13, 42, 73, 104 | 78, 130, 182, 263 |

In general, it is known that reciprocating flows can generate steady streaming, a residual velocity field that remains after averaging over a complete cycle [13, 14]. This phenomenon, often termed “secondary streaming” in the biofluid-mechanical context, distinguishes the time-averaged recirculating vortical motion from the primary axial net flow. Opposing components of this secondary streaming flow may support fluid exchange even when the net transport is small. Exemplary, studies of reciprocating flow through airway bifurcations have demonstrated how streaming can contribute effectively to transport through branching passages [13, 15].

The third ventricle receives CSF from the lateral ventricles through the paired foramina of Monro (FOM) and communicates with the fourth ventricle through the AoS [1]. Flow must pass between the ventricular cavity and these narrow channels. The curved walls and, where present, the interthalamic adhesion (ITA) further constrain its path [16]. These features produce local velocity gradients and may influence whether residual flow predominately circulates within the third ventricle. Earlier computational fluid dynamics (CFD) studies in the third ventricle used subject-specific models to examine aqueductal jets, recirculation and mixing in this region [7, 17]. Those studies focused on the cardiac-driven flow cycles, based on the characteristic heart-beat frequency and pressure pulse. These studies provide the context for the present analysis of residual streaming in a representative human third-ventricle geometry. We vary the parameters stroke volume, frequency and waveform asymmetry, motivated by reported cardiac- and respiration-related CSF motion in human cohorts [4, 11]. We analyse the cycle-averaged velocity fields and preferential flow pathways, and quantify the streaming flux across the AoS as a result of those parameters. The aim is to determine how the organisation of ventricular flow changes and influences the contribution to intra-ventricular streaming.

## Methods

### Geometry and mesh generation

A representative human third-ventricle geometry was derived from the Database BodyParts3D/Anatomography, developed by the Center for Life Science (DBCLS) in Japan [18]. BodyParts3D provides a standard anatomical reference model for simulation studies rather than a population-averaged geometry [18]. It was transferred into Fusion 360 to generate a computational fluid domain that contains the third-ventricle body, the FOM, and the AoS. The reconstructed fluid-domain volume was approximately 1.25 mL, which is within the range of adult third-ventricle volumes reported by Hernández-Cortés et al. [19]. The inferior end of the AoS was extended further into a circular pipe with small taper to ensure smooth entrance into the aqueduct when the flow is rostral. The equivalent diameter of the inferior end with a cross-sectional area of *A*_Aq_ = 2.86 mm^2^ results to *D*_Aq_ = 1.9 mm. The cross-sectional area lies within the range reported in normal subjects [20]. The combined area of the two FOM inlets was *A*_in_ = 11.24 mm^2^.

The processed geometry was imported into ANSYS Meshing. An unstructured tetrahedral mesh was generated using a global element size of 0.09 mm, with local refinement around the FOM, the AoS and strongly curved regions down to 0.533 *µ*m, containing approximately 4.84 million elements. A mesh-independence study was carried out and showed sufficient resolution for all simulated parameters.

### Fluid model and numerical method

CSF was modeled as an incompressible Newtonian fluid with density *ρ* = 1000 kg m^−3^ and kinematic viscosity *ν* = 1.0 *×* 10^−6^ m^2^ s^−1^ [21, 22]. The transient incompressible Navier–Stokes equations were solved in ANSYS Fluent using a pressure-based solver and a laminar formulation. The walls were rigid and no-slip, consistent with previous ventricular CFD modelling [22].

### Boundary conditions and simulation settings

Idealised periodic waveforms of the flow (period *T*) through the cerebral aqueduct were adopted as a first-order approximation to the successive flow pulses driven by the cardiac and breathing cycles. Common to all simulated waveforms is that they have zero net-flow, focusing only on the reciprocating component of CSF transport and its streaming. Realization of this flow profile within the CFD code was enforced by setting time-varying inflow boundary condition at the FOM following the imposed profile with area-equivalent peak value in each of the two FOM. The caudal end of the AoS was then chosen as zero pressure outlet condition. The combination of a prescribed pulsatile velocity and a reference-pressure boundary has precedent in ventricular CFD [7].

The flow evolution was simulated for three complete cycles, while for all simulations the flow in the AoS remains strictly in the laminar regime with a Reynolds-number of *Re* = *U*_Aq,peak_*D*_Aq_*/ν* < 360, defined with the cerebral aqueduct diameter and the peak cross-sectional average velocity. The Womersley number characterises the relative importance of unsteady inertial effects and viscous diffusion in the oscillatory CSF flow [23] as 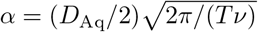. Herein, the maximum reached is 2.4, which is in the intermediate regime, where unsteady inertial effects begin to compete with viscous diffusion. The solver time step was adjusted such that the Courant–Friedrichs–Lewy (CFL) number was always below 0.2, estimated using *C* = *U*_peak_Δ*t/*Δ*x*, with Δ*x* = 0.09 mm as the representative mesh length [24]. This ensures sufficient spatio-temporal resolution for the laminar flow situation in the ventricle.

### Types of waveforms

The flow waveform in the aqueduct was implemented in three different Fourier-type expansions: a pure sinusoidal waveform (1), a phase-modulated waveform (2), and an amplitude modulated waveform (3)

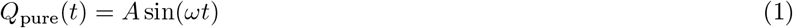

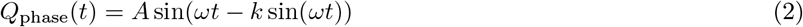

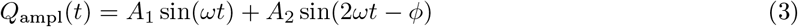

where *A* is the peak aqueductal flow rate and *T* = 2*π/ω* is the oscillation period. In Eq (3), *A*_1_ and *A*_2_ are the first- and second-harmonic flow-rate amplitude coefficients. For the asymmetric waveform prescribed for Fig 2, *A*_1_ = 0.42 mL s^−1^, *A*_2_ = 0.17 mL s^−1^, *ϕ* = 1.0 rad and *T* = 4 s. The time origin is shifted by evaluating Eq (3) at *t* + *t*_0_, with *t*_0_ = 0.1392 s, so that the cycle begins at the zero crossing into the positive half-wave. Positive flow denotes caudal flow towards the fourth ventricle, whereas negative velocity denotes rostral flow towards the third ventricle. For the imposed waveform, the positive stroke volume is defined as

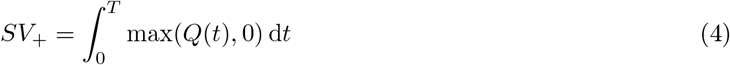

This is the one-way tidal displacement of the fluid bulk through the AoS in cranio-caudal direction, counterbalanced by an equal amount of *SV*_−_ after flow reversal.

The parameter *k* in (2) controls the waveform skewness to replicate timing asymmetry between acceleration and deceleration phase. A parameter *k* = −0.5 leads to a (*T*_*rel*_*≈* 2/3T) which is typical for a cardiac cycle as reported in Beggs et al. [25], driven by a rapid systolic acceleration phase, followed by a slower, prolonged diastolic deceleration (*~* 60%–65% of the cycle) [7, 26–28]. The waveform represents a monotonic phase modulation that shifts the temporal timing of the velocity peaks while preserving their magnitudes (*±A*) and period (*T*). Because *ψ*(*T − t*) = 2*π − ψ*(*t*), the waveform is strictly odd-symmetric about the half-period. The third function combines temporal asymmetry and asymmetry in the peaks. It was generated from a curve fit to a measured waveform from Dreha-Kulaczewski et al. [11], which replicates the rapid, steep cranio-caudal (inspiratory) surge lasting 40% of the cycle, followed by a prolonged, blunted caudo-cranial (expiratory) return lasting 60% of the cycle. The variation of those waveform allows to conclude on the streaming effects when similar fundamental temporal and amplitude skewness exist under resting conditions (cardiac dominated) compared to forced breathing (breathing dominated).

The amplitudes of the waveform functions are scaled in the parametric study such that we can compare the results at comparable aqueductal peak Reynolds-numbers while varying stroke volume and period *T*. The cardiac-frequency group used *T* = 1 s and target aqueductal stroke volumes of 6, 20, 35 and 50 *µ*L to explore the range of reported values for adult normal humans at resting conditions Spijkerman et al. [5]. The longer-period group used target aqueductal stroke volumes of 150, 250, 350 and 500 *µ*L as an exploratory test of the amplitude range reported in [29].

The resulting peak aqueductal velocities are in the range of healthy volunteers, e.g. Lee et al. [30] reported group-mean peak systolic velocities of 3.30–4.08 cm s^−1^, depending on section location and background correction. Sartoretti et al. [31] reported group means of 8.10–10.82 cm s^−1^ for peak velocity within the imaging region of interest, depending on age and sex. Subject-specific CFD by Kurtcuoglu et al. [7] gave local maxima of 16.7 cm s^−1^ caudally and 14.7 cm s^−1^ cranially in the narrowest aqueductal region. These values provide context for the chosen amplitudes. All parameter variations are given in Table 1.

### Post-processing and flow metrics

In general, the Lagrangian mass flux in oscillating flows is the sum of the time-averaged Eulerian streaming velocity **Û**_*s*_ and the kinematic Stokes drift. In the current flow situation with separated jet-sink regime as shown later, macroscopic transport is governed by topological asymmetry between directional jet ejection and omnidirectional sink suction, alongside advective Reynolds stresses, characterised by the Eulerian streaming field. The latter is analysed in detail using streamline patterns and contour plots in the midsagittal plane. The component of residual streaming flux across an arbitrary cross-sectional plane *A* through the volume is calculated as follows (see also Wanigasekara et al. [15]):

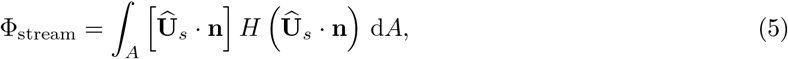

where *H* is the Heaviside step function and **n** the vector normal to the plane. To evaluate the cross-ventricular streaming flux, two cross-sections were chosen through the AoS, at the superior and inferior position, see Fig 1. The unit normal vector **n** in Eq (5) then points in the selected planes in caudal direction. Φ_stream_ represents the positive streaming flux (caudal), which must be balanced by equal amount of negative flux (rostral) to fulfill mass conservation. The streaming ratio was then defined as

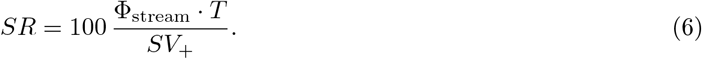

*SR* measured in the AoS represents the percentage of the reciprocating stroke volume that is rectified into continuous bidirectional exchange between the 3rd and 4th ventricle.

**Fig 1.**
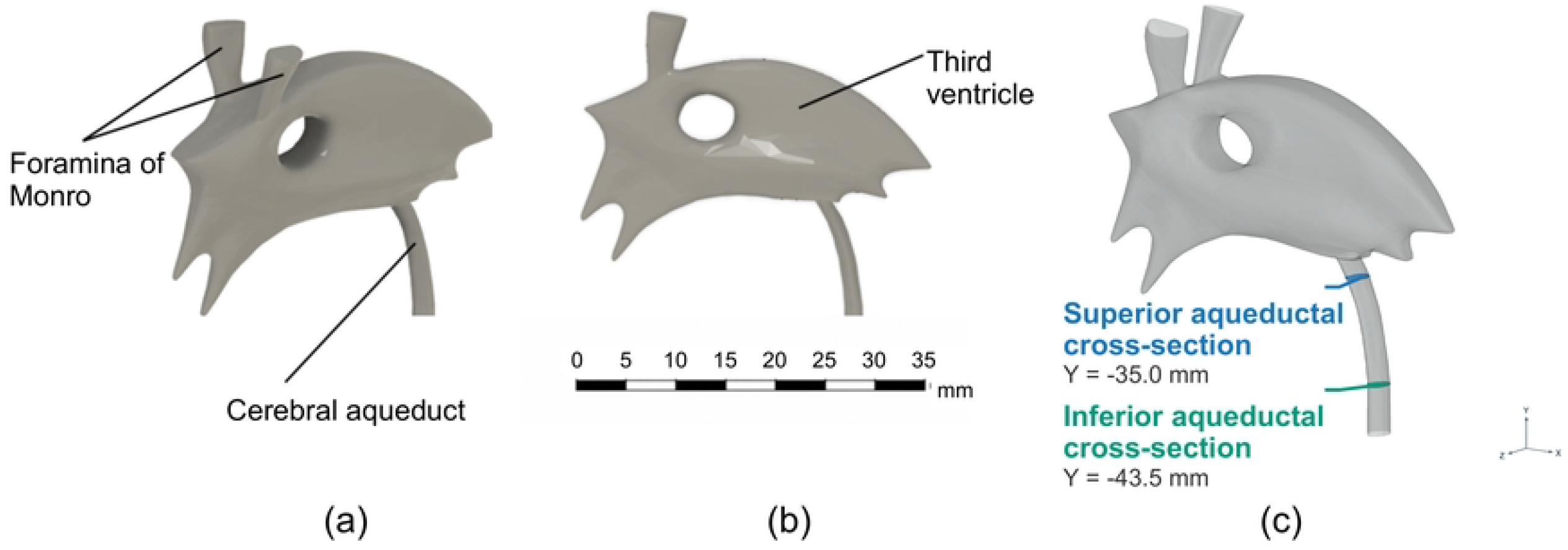
Reconstructed third-ventricle geometry used for CFD simulation, derived from BodyParts3D/Anatomography (Database Center for Life Science, Japan; [18]). (a) Three-dimensional view of the processed fluid domain. (b) Lateral view. The vertical axis of the coordinate system is aligned with the axis of the inferior end of the cerebral aqueduct. (c) Locations of the superior and inferior aqueductal cross-sections used for flow analysis.

**Fig 2.**
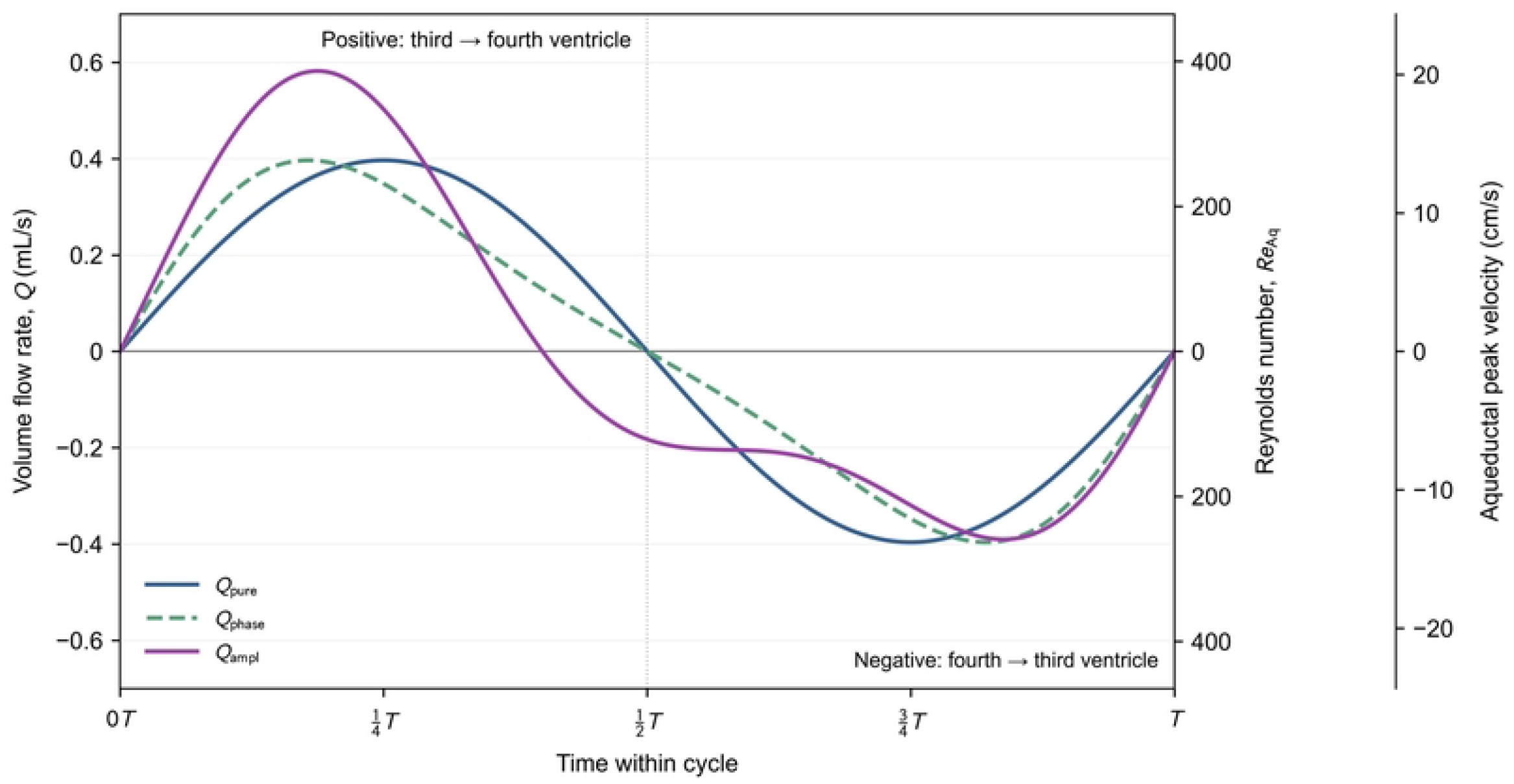
Exemplary plots of the implemented aqueductal flow waveforms at a fixed peak aqueductal Reynolds number of. Re = 263. The right-hand axis shows the aqueduct peak velocity and corresponding Reynolds number. The plots represent one combination from the parametric variation study.

## Results

### Instantaneous bidirectional flow over the cycle

The instantaneous velocity fields were compared at nine equally spaced phases of the final simulated cycle (Fig 3). Flow is displayed by means of sectional streamlines representing 2D planar projections of a 3D flow field in this plane. The normalised phase was defined as *t*\* = (*t − t*_0_)*/T*, where *t*_0_ is the start of the analysed cycle and *T* is the oscillation period. Thus, *t*\* = 0 and *t*\* = 1 represent equivalent phases in successive cycles. Under the prescribed waveform, 0 *< t*\* *<* 1*/*2 corresponds to caudal flow from the FOM towards the AoS, whereas 1*/*2 *< t*\* *<* 1 corresponds to rostral/cephalad reverse flow. The phases *t*\* = 0, 1*/*2, and 1 are associated with flow reversals, at which the imposed inlet velocity approaches zero.

**Fig 3.**
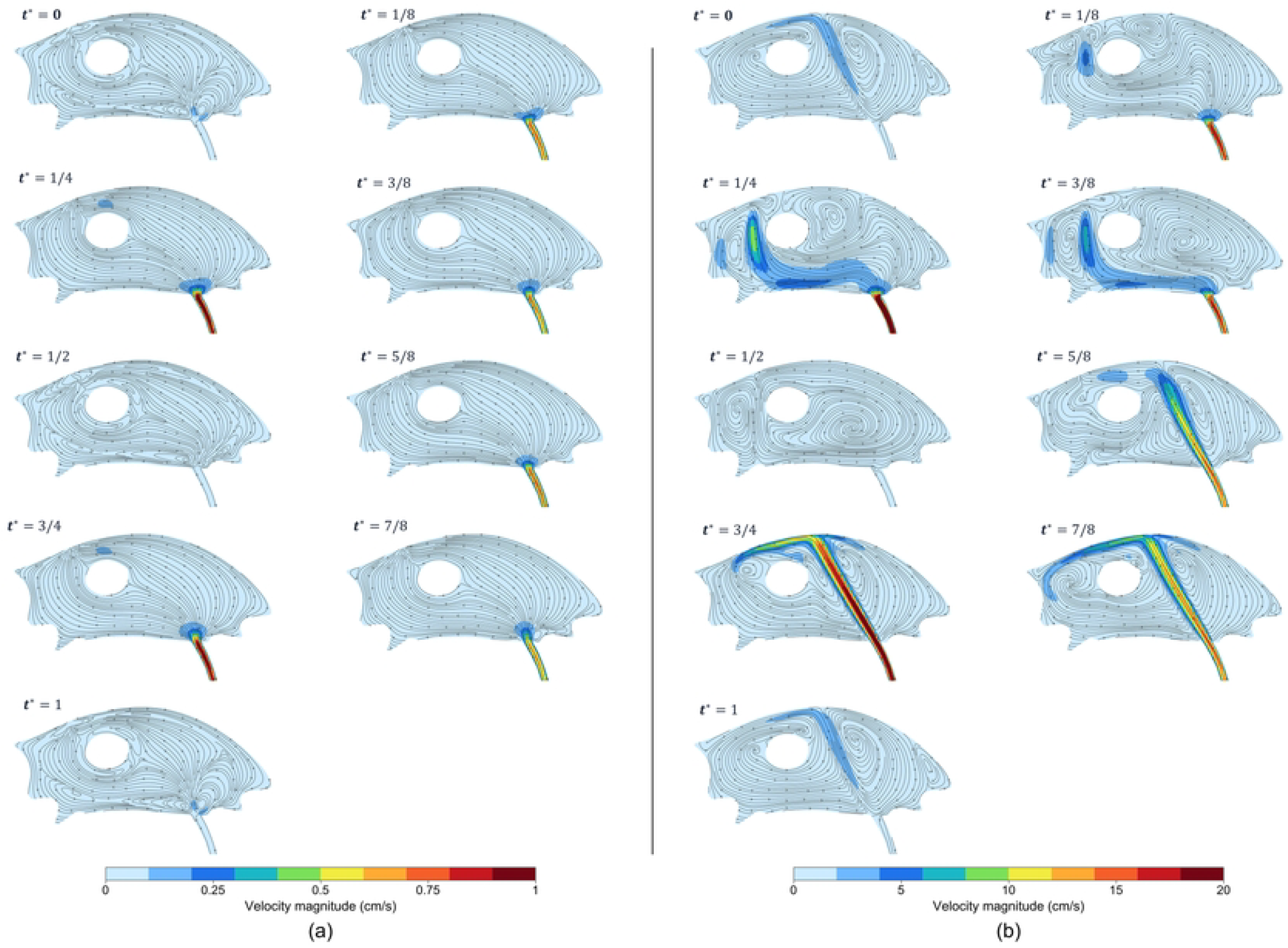
Instantaneous midsagittal velocity magnitude and in-plane streamlines in one wave-cycle (waveform *pure*), separated by. Δ*t*\* = 1*/*8. (a) *Re*_Aq,peak_ = 13, *T* = 1 s; (b) *Re*_Aq,peak_ = 263, *T* = 4 s.

For the lower-amplitude *T* = 1 s case, the highest speeds were confined mainly to the aqueduct, while most of the ventricular cavity remained at low velocity. At the peak caudal and rostral phases, *t*\* = 1*/*4 and 3*/*4, the broad streamline paths were similar, indicating the tendency of the low-Reynolds number flow to preserve a near time-reversal feature.

For the higher-amplitude *T* = 4 s case, the caudal flow followed the left side of the ITA towards the AoS. During the rostral phase, a concentrated aqueductal jet entered the cavity along the right side of the ITA. The two half-cycles therefore followed different paths, with local recirculation around the jets.

### Streaming flow organisation

The time-averaged midsagittal streaming velocity field is shown in Fig 4 by means of sectional streamlines and color-coded contours of magnitude of in-plane velocity. In the low-Re case, velocities were small across most of the cavity and only higher near the FOM and AoS. Inside the ventricle, a broad clock-wise circulation occurs around the ITA. A pair of counter-rotating vortices is observed at the entrance of the AoS, indicating the jet penetration zone. In the small diameter aqueduct at 6 *µ*L the plug of fluid travels a distance of *L* (*L* = *SV/A*_*Aq*_) *≈* 2.1*mm*, leading to *L/D*_*Aq*_ *<* 4 which is below the limit of *L/D ≈* 4 given by Gharib [32] for the formation and pinch-off of a leading vortex ring; the fluid merely wobbles at the exit and is sucked right back in upon reversal. The results prove that that vortex has not yet detached from the ventricle walls in the cycle. Across the rest of the parametric sweep at *SV* = 20 *µ*L, *L/D*_Aq_ *≈* 3.7 approaches the pinch-off limit, and for *SV* = 35–500 *µ*L, *L/D*_Aq_ spans 6.4 to 92, far exceeding *L/D ≈* 4.

**Fig 4.**
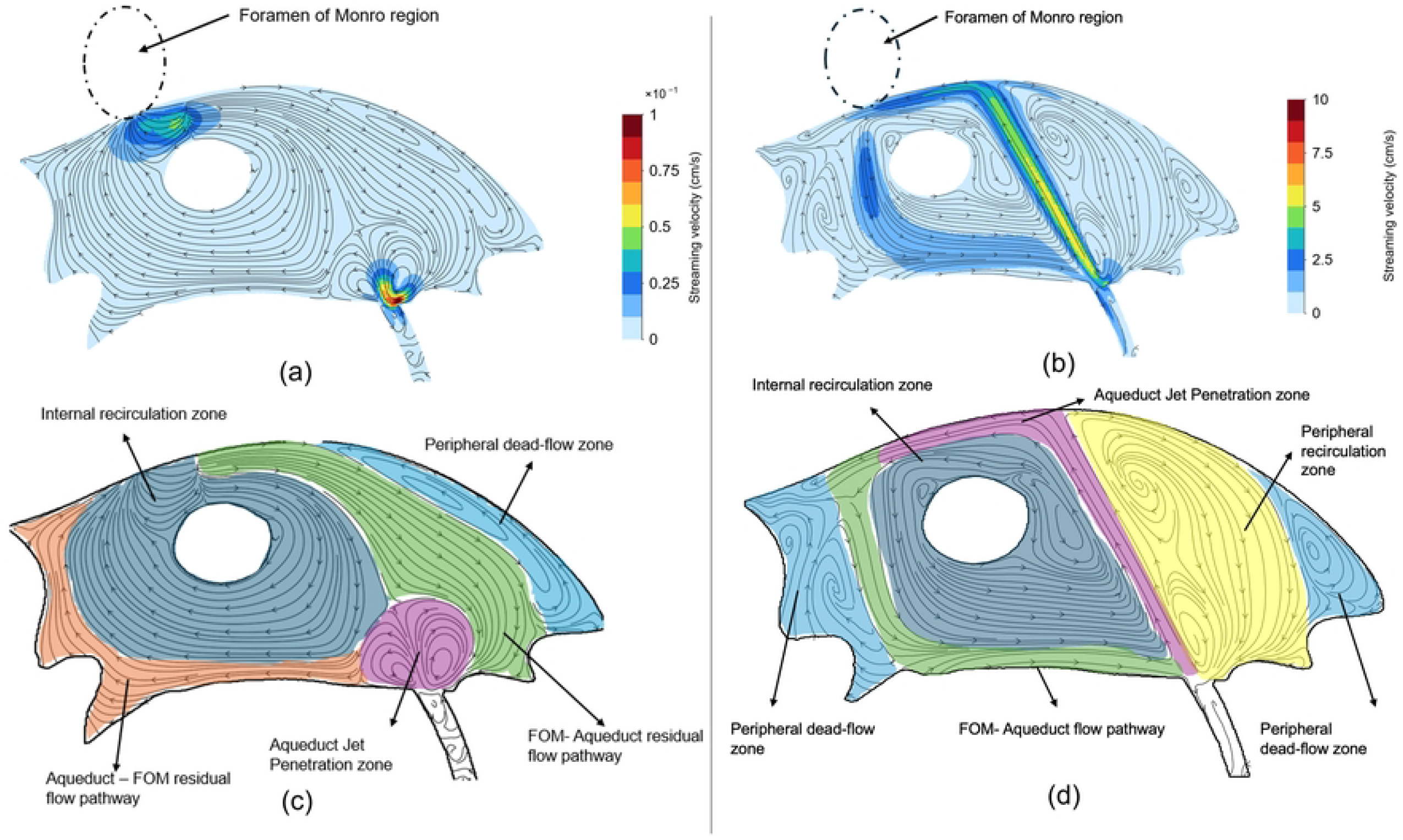
Streaming flow organisation in the midsagittal plane for low and high. *Re*_Aq,peak_. Case a,c (waveform *pure, T* = 1 s, *Re*_Aq,peak_ = 13) compared to case b,d (waveform *pure, T* = 4 s, *Re*_Aq,peak_ = 263). Dashed ellipses mark the approximate projected FOM regions; panels (c,d) show qualitative annotations of pathways and recirculating regions.

In the high-Re case, two bands of elevated mean velocity pathway exist, one extended rostrally from the AoS along the right side of the ITA and towards the superior ventricular region. The return path passed caudally around the left and lower parts of the cavity towards the AoS. Both pathways represent the passage of the strong jets, now penetrating deep and interacting with the opposite walls. The central and peripheral recirculating regions were separated by these stronger mean-flow pathways. Between both counter-acting pathways, a larger recirculation region rotating counter clock-wise is formed around the ITA. This region of coherent rotating motion covers half of the dimension of the ventricle and is energized by the shearing motion of both pathways high-velocity regions. Thus, with increasing Re-numbers, preferential pathways are formed which drive the internal recirculating region to switch sign of rotation and form a strong fluidic “flywheel” inside the ventricle.

Fig 5 compares the midsagittal streaming flow fields for the different waveforms. This shows, that the dominating parameter for the flow pathways is rather the peak Reynolds-number than the waveform.

**Fig 5.**
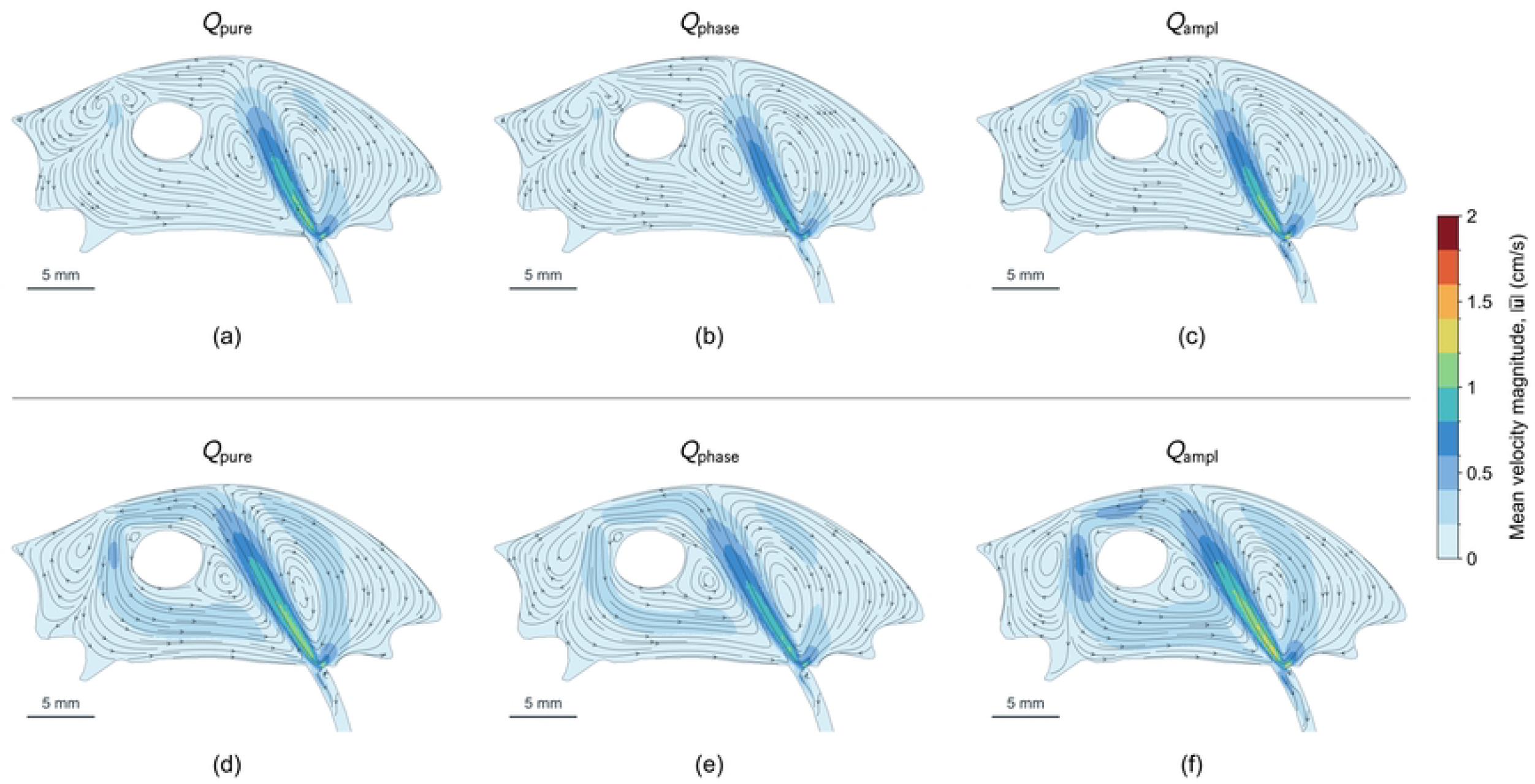
**Streaming flow organisation for different waveforms at similar** *Re*_Aq,peak_ **caudo-cranial; panels (a,b,c)**: *Re*_Aq,peak_ = 73, *T* = 1 s; **panels (d,e,f)**: *Re*_Aq,peak_ = 78, *T* = 4 s.

### Streaming across the aqueduct

While the streaming flow pattern in the third ventricle illustrates local jet formation and internal recirculation, evaluating the residual streaming flux across the AoS provides the metric of cross-ventricular transport. We selected the superior and inferior aqueductal cross-sections as shown in Fig 1(c) for further analysis of the structure of the streaming pattern. Again, flow is displayed by means of sectional streamlines representing 2D planar projections of the 3D flow field in this plane.

Both aqueductal sections demonstrate the pathway regions of positive and negative streaming normal to the plane of the cross-section (Fig 6). At the superior cross-section, the residual velocity field exhibits a pronounced asymmetric, bidirectional organization: net rostral streaming is concentrated within the central core, whereas net caudal streaming is localized along the inner wall of the curved channel. This is observed for the low and the high Reynolds-number case in similar pattern, but two orders of magnitude difference in the velocities. This pattern shows a sign-reversal when moving further caudal in the aqueduct. Then, the core region shows pronounced caudal flow while the region around the inner wall of the curved channel has rostral flow. This undulation of the streaming pathway along the longitudinal axis is interpreted as the result of the secondary circulation, seen by the formation of Dean-type vortices. In curved passages, the mean motion can include both axial streaming and secondary circulation, whose spatial organisation varies along the channel. Such behaviour has been demonstrated in studies of oscillatory flow through curved tubes [33, 34].

**Fig 6.**
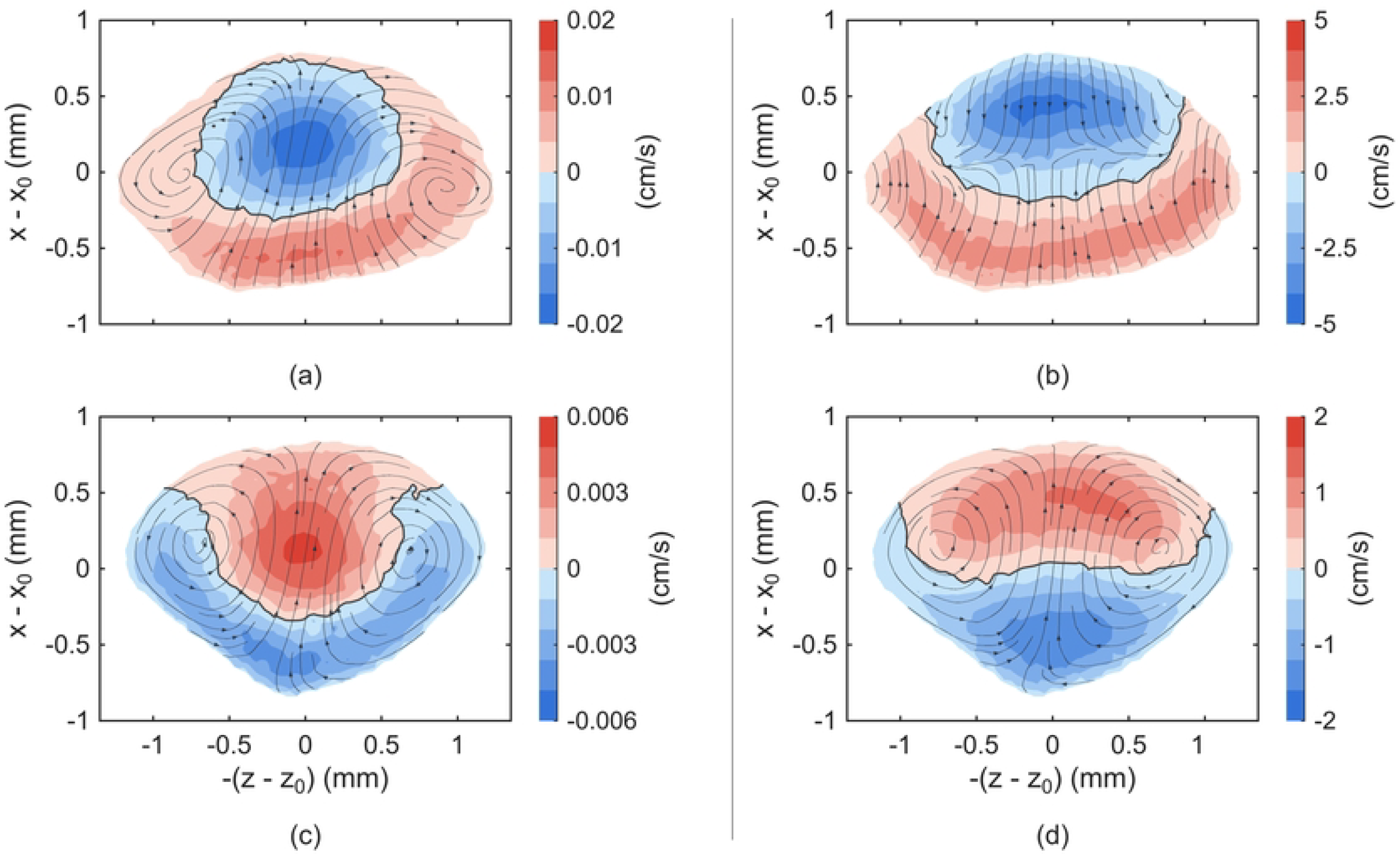
Contours of out-of-plane streaming velocity Û_*s*_ · n at (a,b) the superior and (c,d) the inferior aqueductal cross-section, waveform. *pure*. Overlaid are cross-sectional streamlines of the streaming field. Left column: *Re*_Aq,peak_ = 13, *T* = 1 s; right column *Re*_Aq,peak_ = 263, *T* = 4 s. Red denotes caudal flow and blue rostral flow.

When analyzing the quantitative values of *SR* we found the dominant parameter to be the peak aqueductal Reynolds number as shown in Fig 7 for both aqueductal cross-sections. A good approximation of the measured results is described by the hyperbolic saturation model *SR* = *ARe/*(*B* + *Re*), with the coefficients given in the figure. Note that the Re-values for the asymmetric waveform are taken as the absolute peak aqueductal Re-number in caudo-cranial direction for correct reference.

**Fig 7.**
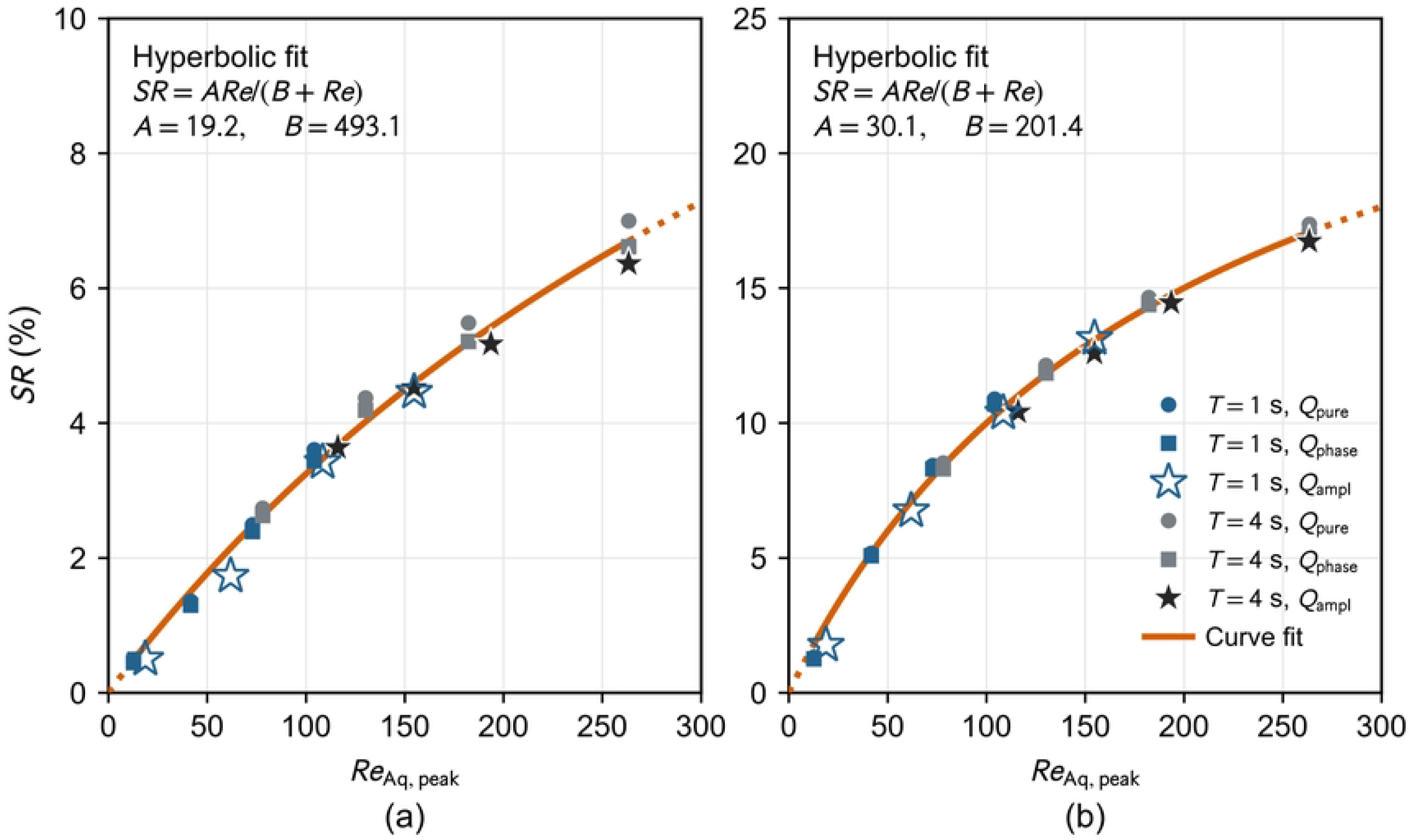
Streaming ratio as function of peak aqueductal Reynolds number (a) inferior and (b) superior aqueductal cross-section. Data fitted to a hyperbolic saturation model, *SR* = *ARe/*(*B* + *Re*); fit goodness statistics are *R*^2^ = 0.990, *RMSE* = 0.194 (inferior) and *R*^2^ = 0.994, *RMSE* = 0.371 (superior).

## Discussion

This study compared the CSF flow streaming pattern in the third ventricle in a parametric study of varying stroke volumes and waveforms, simulating the broader range of observed parameters in physiological CSF flow studies under normal and forced breathing conditions. The parameter sweep extends previous studies of third-ventricle flow and mixing [7, 17] by comparing period, amplitude and waveform shape in one geometry. While at resting conditions the CSF pulsating motions are dominated by the cardiac cycle, forced breathing changes that to slower but stronger fluid sloshes. The results show, that the forced breathing waveform produced greater velocity magnitude and streaming exchange, together with clearer caudal and rostral preferential pathways. Similar qualitative streaming pattern however could also be observed for high-frequency cardiac-driven waveforms at the upper end of stroke volumes. A comparison of the flow patterns at the same peak Reynolds-number documented this matching. Although these regimes operate at distinct Womersley numbers (*α* varying by a factor of two), the resulting steady streaming field is effectively not changed. Furthermore, the parametric change of the waveform also did not change the global streaming pattern, except from regions close the FOM and AoS. Therefore it is concluded that the waveform in the given range of parameters has not much of an effect.

The overall dominating factor in the streaming pattern is the peak aqueductal jet momentum (*Re*_Aq, peak_). The formation of preferential pathways with increasing Reynolds-number is related to deeper penetration of the jets from the FOM and AoS in the ventricle and their interaction with the walls. The laterally parallel but separated caudal and rostral pathways have trapped an inner recirculating region around the ITA, which is circulating as a coherent fluid bulk counter-clockwise, forming a fluidic “flywheel”. This region is continuously re-energized by the alternating jets passing by and shearing the fluid.

The streaming ratio measured in the superior aqueduct increased from 1.3% to 17.3% with increasing Re-number from 13 to 263, indicating that both the strength of oscillatory exchange and the organisation of the residual streaming flow pattern enhanced cross-ventricular transport. The observed trend fits to high statistical goodness to a hyperbolic saturation function. For all cases with waveform *pure*, the streaming values are slightly higher than the fitting curve, while the waveform *ampl* showed somewhat smaller values. Those differences are observed to be less than 5% of the corresponding fitted trend. Moving further caudal along the longitudinal axis of the aqueduct to the inferior end the results show reduced streaming fluxes by approximately half. Nevertheless, this flux represents a considerable amount of steady cross-ventricular transport.

It should be noted that the observed streaming ratio is a hydrodynamic descriptor and cannot be interpreted directly as a measure of physiological clearance efficiency, which should be investigated in future work. However, our findings may suggest that streaming is relevant as a further mechanism that could support metabolite clearance [35], a mechanism not previously studied in this context.

## Conclusion

This study compared cerebrospinal fluid (CSF) motion in one representative human third-ventricle geometry under reciprocating, bi-phasic flow conditions representative of the pulse waves induced by the cardiac and respiratory cycles during resting and forced breathing conditions. The latter being an indicator for sleep-related forcing of stronger bulk CSF displacement; compared to previous studies, we focused on the residual streaming patterns, a feature that indicates transport and mixing in an otherwise net-zero reciprocating flow. The simulated conditions reproduced aqueductal peak velocities well within the range of reported quantities for healthy volunteers (1–14 cm/s). Towards the lower band of stroke volumes(*SV* = *O*(10)*µL*), the resulting streaming flow features resemble an approximate time-reversal flow pattern. Increasing the stroke-volume and/or the frequency separates those flow passages into opposite directed and laterally separated jets entering the ventricle in an alternating manner driven with the cyclic flow conditions. Those jets with a typical Reynolds number of *Re* = *O*(100) stem in the cranio-caudal cycle from the FOM, while - after flow reversal - the jet entering from the AoS dominates the flow pathway. The preferential pathways are therefore related to the alternating source-sink features of the flow at the FOM and the AoS and the geometrical features of the ventricle with its internal structures and position / orientation of the connecting conduits of FOM and AoS. As a result, for the given geometry a larger inner recirculation zone is formed and trapped between both high-velocity pathways of which circulation is re-energized by the shear layer along the jets with each wave. This “flywheel” contributes to the cross-ventricular mixing of the fluid entering from both conduits.

Quantitative assessment of streaming transport across the AoS represented a considerable proportion of the cross-ventricular fluid exchange. Ultimately, our findings demonstrate that this permanent net-transport due to streaming is governed primarily by the peak aqueductal jet Reynolds number (*Re*_peak_). This exchange is best characterized by a hyperbolic saturation function, *SR ~ Re/*(*B* + *C · Re*). This relationship captures the physical transition between two governing regimes: at low inertial flow, streaming grows relative to the oscillating flow approximately linearly with *Re*, the entering jet expands rather freely without impingement, yielding a near-linear conversion of peak advective momentum into residual circulation. At elevated Reynolds numbers however, the jet interaction with the structural boundaries of the third ventricle dissipate excess kinetic energy. The ventricular anatomy therefore constrains residual streaming against escalating inertial fluid sloshing.

Our conclusions are related to the findings for the chosen specific geometry of the 3rd ventricle, including a prominent interthalamic adhesion (ITA). The interaction of the flow with this internal boundary is a separate subject to be studied in further research requesting another parametric study as size and position of the ITA vary considerably in human population. Previous work had already investigated the influence of ITA on instantaneous flow dynamics in the 3rd ventricle, see e.g. [36, 37]. Furthermore, our chosen waveforms are approximations of reported profiles from literature and are applied as isolated dominant 1Hz or 0.25 Hz frequencies. Although aqueductal flow at *Re* = *O*(100) is inertia-dominated, isolating the 0.25 Hz respiratory wave from the 1.0 Hz cardiac ripple is considered as a reasonable approximation for forced breathing conditions. The high-volume respiratory jet carries over 95% of the driving kinetic energy. While cardiac ripples may slightly agitate the edges of the entering jet, the global streaming pattern and net cross-ventricular transport are governed almost entirely by the dominant respiratory flush.

## Acknowledgments

This paper is dedicated to the memory of Luke McTavish. The authors thank Lady Dorit Young of Dartington for her interest and inspiration to further investigate flow in the human ventricles as part of the complex metabolic system of the brain.

## Data availability

The data that support the findings of this study are available from the corresponding author upon reasonable request.

## Declaration of generative AI and AI-assisted technologies in the manuscript preparation process

During the preparation of this work, the authors used OpenAI ChatGPT/Codex to assist with language editing, formatting, and manuscript organisation. After using this tool, the authors reviewed and edited the content as needed and take full responsibility for the content of the publication.

## Appendix A. Numerical sensitivity and cycle repeatability

Numerical sensitivity was assessed for the largest pure-sinusoidal case (*k* = 0, *T* = 4 s, target *SV* = 500 *µ*L, *Re*_Aq,peak_ *≈* 263). The baseline and refined meshes contained 4,843,127 and 8,497,703 tetrahedra. Time steps of 0.500, 0.250 and 0.125 ms were compared over 8–12 s from the same 8 s restart. Mesh comparisons used the third cycle; repeatability was checked between cycles 2 and 3. *SR* was normalised by *SV*_+_.

**Table A.1.**
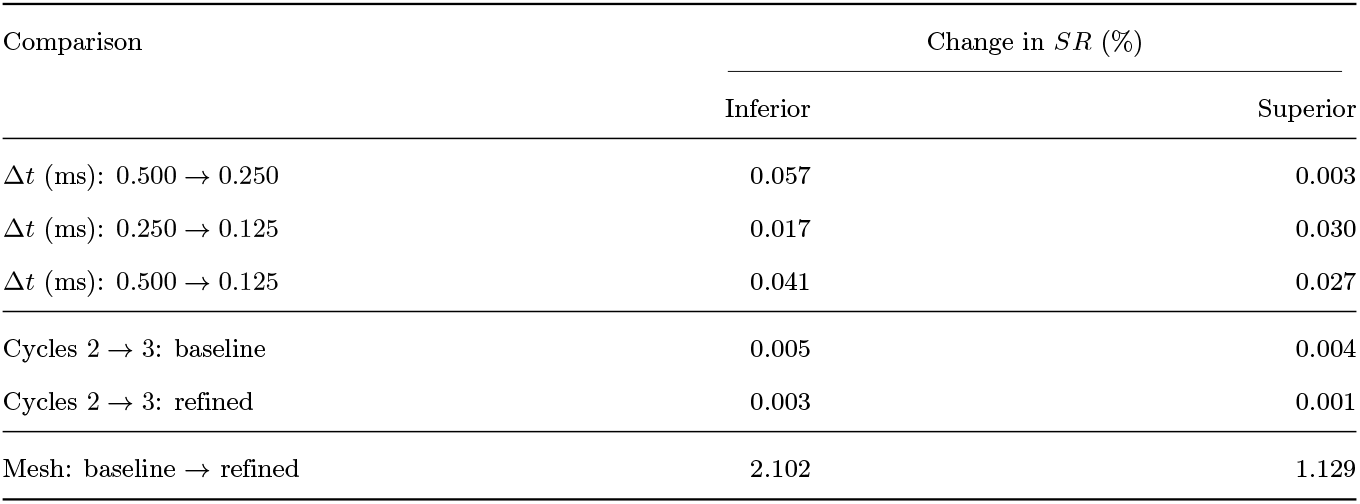
Time-step, cycle-repeatability and SR-based mesh comparisons.

| Comparison | Change in *SR* (%) |  |
|---|---|---|
|  | Inferior | Superior |
|---|---|---|
| Δ*t* (ms): 0.500 → 0.250 | 0.057 | 0.003 |
| Δ*t* (ms): 0.250 → 0.125 | 0.017 | 0.030 |
| Δ*t* (ms): 0.500 → 0.125 | 0.041 | 0.027 |
| Cycles 2 → 3: baseline | 0.005 | 0.004 |
| Cycles 2 → 3: refined | 0.003 | 0.001 |
| Mesh: baseline → refined | 2.102 | 1.129 |

The time-step and cycle-repeatability checks were completed and satisfied the 2% criterion for *SR*. Mesh testing was also completed: refinement changed *SR* by 2.102% at the inferior cross-section and 1.129% at the superior cross-section.

## References

1. Sakka L, Coll G, Chazal J. Anatomy and physiology of cerebrospinal fluid. European Annals of Otorhinolaryngology, Head and Neck Diseases. 2011;128(6):309–16. doi:10.1016/j.anorl.2011.03.002.

2. Iliff JJ, Wang M, Liao Y, Plogg BA, Peng W, Gundersen GA, et al. A paravascular pathway facilitates CSF flow through the brain parenchyma and the clearance of interstitial solutes, including amyloid β. Science Translational Medicine. 2012;4(147):147ra111. doi:10.1126/scitranslmed.3003748.

3. Yamada S, Miyazaki M, Yamashita Y, Ouyang C, Yui M, Nakahashi M, et al. Influence of respiration on cerebrospinal fluid movement using magnetic resonance spin labeling. Fluids and Barriers of the CNS. 2013;10:36. doi:10.1186/2045-8118-10-36.

4. Chen L, Beckett A, Verma A, Feinberg DA. Dynamics of respiratory and cardiac CSF motion revealed with real-time simultaneous multi-slice EPI velocity phase contrast imaging. NeuroImage. 2015;122:281–7. doi:10.1016/j.neuroimage.2015.07.073.

5. Spijkerman JM, Geurts LJ, Siero JCW, Hendrikse J, Luijten PR, Zwanenburg JJM. Phase Contrast MRI Measurements of Net Cerebrospinal Fluid Flow Through the Cerebral Aqueduct Are Confounded by Respiration. Journal of Magnetic Resonance Imaging. 2019;49(2):433–44. doi:10.1002/jmri.26181.

6. Abbey P, Singh P, Khandelwal N, Mukherjee KK. Shunt surgery effects on cerebrospinal fluid flow across the aqueduct of Sylvius in patients with communicating hydrocephalus. Journal of Clinical Neuroscience. 2009;16(4):514–8. doi:10.1016/j.jocn.2008.05.009.

7. Kurtcuoglu V, Soellinger M, Summers P, Boomsma K, Poulikakos D, Boesiger P, et al. Computational investigation of subject-specific cerebrospinal fluid flow in the third ventricle and aqueduct of Sylvius. Journal of Biomechanics. 2007;40(6):1235–45. doi:10.1016/j.jbiomech.2006.05.031.

8. Dreha-Kulaczewski S, Konopka M, Joseph AA, Kollmeier J, Merboldt KD, Ludwig HC, et al. Respiration and the watershed of spinal CSF flow in humans. Scientific Reports. 2018;8(1):5594. doi:10.1038/s41598-018-23908-z.

9. Yatsushiro S, Sunohara S, Tokushima T, Takizawa K, Matsumae M, Atsumi H, et al. Characterization of Cardiac- and Respiratory-driven Cerebrospinal Fluid Motions Using a Correlation Mapping Technique Based on Asynchronous Two-dimensional Phase Contrast MR Imaging. Magnetic Resonance in Medical Sciences. 2021;20(4):385–95. doi:10.2463/mrms.mp.2020-0085.

10. Vijayakrishnan Nair V, Kish BR, Inglis B, Yang HCS, Wright AM, Wu YC, et al. Human CSF movement influenced by vascular low frequency oscillations and respiration. Frontiers in Physiology. 2022;13:940140. doi:10.3389/fphys.2022.940140.

11. Dreha-Kulaczewski S, Joseph AA, Merboldt KD, Ludwig HC, Gärtner J, Frahm J. Inspiration is the major regulator of human CSF flow. The Journal of Neuroscience. 2015;35(6):2485–91. doi:10.1523/JNEUROSCI.3246-14.2015.

12. Elabasy A, Helakari H, Väyrynen T, Rajna Z, Huotari N, Raitamaa L, et al. Sleep Alters the Velocity of Physiological Brain Pulsations in Humans. Advanced Science. 2026;13(19):e03745. doi:10.1002/advs.202503745.

13. Haselton FR, Scherer PW. Flow Visualization of Steady Streaming in Oscillatory Flow Through a Bifurcating Tube. Journal of Fluid Mechanics. 1982;123:315–33. doi:10.1017/S0022112082003085.

14. Khani M, Sass LR, Xing T, Sharp MK, Balédent O, Martin BA. Anthropomorphic Model of Intrathecal Cerebrospinal Fluid Dynamics Within the Spinal Subarachnoid Space: Spinal Cord Nerve Roots Increase Steady-Streaming. Journal of Biomechanical Engineering. 2018;140(8):081012. doi:10.1115/1.4040401.

15. Wanigasekara C, Jacob C, Manasseh R, Leontini J. Mean streaming in reciprocating flow in a double bifurcation. Fluid Dynamics Research. 2024;56(4):045505. doi:10.1088/1873-7005/ad6289.

16. Duque Parra JE, Pava Ripoll Á, Vélez García JF. Interthalamic Adhesion in Humans: A Gray Commissure? Anatomy & Cell Biology. 2022;55(1):109–12. doi:10.5115/acb.21.164.

17. Kurtcuoglu V, Soellinger M, Summers P, Poulikakos D, Boesiger P. Mixing and modes of mass transfer in the third cerebral ventricle: A computational analysis. Journal of Biomechanical Engineering. 2007;129(5):695–702. doi:10.1115/1.2768376.

18. Mitsuhashi N, Fujieda K, Tamura T, Kawamoto S, Takagi T, Okubo K. BodyParts3D: 3D structure database for anatomical concepts. Nucleic Acids Research. 2009;37(Database issue):D782–5. doi:10.1093/nar/gkn613.

19. Hernández-Cortés KS, Pujals-Table AA, Sagaró del Campo-Nelsa M, Montoya-Arquímedes P. Tables of percentiles of the third ventricle according to age and sex associated with brain aging. Revista Mexicana de Neurociencia. 2023;24(5):140–9. doi:10.24875/RMN.23000009.

20. Oner Z, Sagir Kahraman A, Kose E, Oner S, Kavakli A, Cay M, et al. Quantitative evaluation of normal aqueductal cerebrospinal fluid flow using phase-contrast cine MRI according to age and sex. The Anatomical Record. 2017;300(3):549–55. doi:10.1002/ar.23514.

21. Bloomfield IG, Johnston IH, Bilston LE. Effects of proteins, blood cells and glucose on the viscosity of cerebrospinal fluid. Pediatric Neurosurgery. 1998;28(5):246–51. doi:10.1159/000028659.

22. Maeda S, Otani T, Yamada S, Watanabe Y, Yavuz Ilik S, Wada S. Biomechanical effects of hyper-dynamic cerebrospinal fluid flow through the cerebral aqueduct in idiopathic normal pressure hydrocephalus patients. Journal of Biomechanics. 2023;156:111671. doi:10.1016/j.jbiomech.2023.111671.

23. Womersley JR. Method for the calculation of velocity, rate of flow and viscous drag in arteries when the pressure gradient is known. The Journal of Physiology. 1955;127(3):553–63. doi:10.1113/jphysiol.1955.sp005276.

24. Courant R, Friedrichs K, Lewy H. Über die partiellen Differenzengleichungen der mathematischen Physik. Mathematische Annalen. 1928;100:32–74. doi:10.1007/BF01448839.

25. Beggs CB, Shepherd SJ, Cecconi P, Lagana MM. Predicting the Aqueductal Cerebrospinal Fluid Pulse: A Statistical Approach. Applied Sciences. 2019;9(10):2131. doi:10.3390/app9102131.

26. Balédent O, Henry-Feugeas MC, Idy-Peretti I. Cerebrospinal fluid dynamics and relation with blood flow: a magnetic resonance study with semiautomated cerebrospinal fluid segmentation. Investigative Radiology. 2001;36(7):368–77. doi:10.1097/00004424-200107000-00003.

27. Balédent O, Gondry-Jouet C, Meyer ME, De Marco G, Le Gars D, Henry-Feugeas MC, et al. Relationship between cerebrospinal fluid and blood dynamics in healthy volunteers and patients with communicating hydrocephalus. Investigative Radiology. 2004;39(1):45–55. doi:10.1097/01.rli.0000100892.87214.49.

28. Gupta S, Soellinger M, Boesiger P, Poulikakos D, Kurtcuoglu V. Three-dimensional computational modeling of subject-specific cerebrospinal fluid flow in the subarachnoid space. Journal of Biomechanical Engineering. 2009;131(2):021010. doi:10.1115/1.3005171.

29. Vinje V, Ringstad G, Lindstrøm EK, Valnes LM, Rognes ME, Eide PK, et al. Respiratory influence on cerebrospinal fluid flow – a computational study based on long-term intracranial pressure measurements. Scientific Reports. 2019;9:9732. doi:10.1038/s41598-019-46055-5.

30. Lee JH, Lee HK, Kim JK, Kim HJ, Park JK, Choi CG. CSF flow quantification of the cerebral aqueduct in normal volunteers using phase contrast cine MR imaging. Korean Journal of Radiology. 2004;5(2):81–6. doi:10.3348/kjr.2004.5.2.81.

31. Sartoretti T, Wyss M, Sartoretti E, Reischauer C, Hainc N, Graf N, et al. Sex and age dependencies of aqueductal cerebrospinal fluid dynamics parameters in healthy subjects. Frontiers in Aging Neuroscience. 2019;11:199. doi:10.3389/fnagi.2019.00199.

32. Gharib M, Rambod E, Shariff K. A universal time scale for vortex ring formation. Journal of Fluid Mechanics. 1998;360:121–40. doi:10.1017/S0022112097008410.

33. Lyne WH. Unsteady viscous flow in a curved pipe. Journal of Fluid Mechanics. 1971;45(1):13–31. doi:10.1017/S0022112071002970.

34. Sumida M. Steady streaming motion in entrance region of curved tubes during oscillatory flow. Journal of Fluid Flow, Heat and Mass Transfer. 2017;4:36–46. doi:10.11159/jffhmt.2017.005.

35. Xie L, Kang H, Xu Q, Chen MJ, Liao Y, Thiyagarajan M, et al. Sleep drives metabolite clearance from the adult brain. Science. 2013;342(6156):373–7. doi:10.1126/science.1241224.

36. Cheng S, Tan K, Bilston LE. The effects of the interthalamic adhesion position on cerebrospinal fluid dynamics in the cerebral ventricles. Journal of Biomechanics. 2010;43(3):579–82. doi:10.1016/j.jbiomech.2009.10.002.

37. Yamada S, Okada K, Ito H, Iseki C, Yamanaka T, Tanikawa M, et al. Interthalamic Adhesion as a Potential Structural Regulator of Cerebrospinal Fluid Dynamics in the Third Ventricle. Aging and Disease. 2025. Advance online publication. doi:10.14336/AD.2025.1352.

